# Evolutionary analysis of SARS-CoV-2 spike protein for its different clades

**DOI:** 10.1101/2020.11.24.396671

**Authors:** Matías J. Pereson, Diego M. Flichman, Alfredo P. Martínez, Patricia Baré, Gabriel H. Garcia, Federico A. DI Lello

## Abstract

**Objective:** The spike protein of SARS-CoV-2 has become the main target for antiviral and vaccine development. Despite its relevance, there is scarce information about its evolutionary traces. The aim of this study was to investigate the diversification patterns of the spike for each clade of SARS-CoV-2 through different approaches.

**Methods:** Two thousand and one hundred sequences representing the seven clades of the SARS-CoV-2 were included. Patterns of genetic diversifications and nucleotide evolutionary rate were estimated for the spike genomic region.

**Results:** The haplotype networks showed a star shape, where multiple haplotypes with few nucleotide differences diverge from a common ancestor. Four hundred seventy nine different haplotypes were defined in the seven analyzed clades. The main haplotype, named Hap-1, was the most frequent for clades G (54%), GH (54%), and GR (56%) and a different haplotype (named Hap-252) was the most important for clades L (63.3%), O (39.7%), S (51.7%), and V (70%). The evolutionary rate for the spike protein was estimated as 1.08 x 10^−3^ nucleotide substitutions/site/year. Moreover, the nucleotide evolutionary rate after nine months of pandemic was similar for each clade.

**Conclusions:** In conclusion, the present evolutionary analysis is relevant since the spike protein of SARS-CoV-2 is the target for most therapeutic candidates; besides, changes in this protein could have consequences on viral transmission, response to antivirals and efficacy of vaccines. Moreover, the evolutionary characterization of clades improves knowledge of SARS-CoV-2 and deserves to be assessed in more detail since re-infection by different phylogenetic clades has been reported.

## 1. Introduction

In December 2019, the Severe Acute Respiratory Syndrome Coronavirus 2 (SARS-CoV-2) emerged and shocked the entire world (Zhu et al. 2020). After 10 months of worldwide circulation, more than 55 million cases and 1.4 million deaths have been reported globally (WHO, 2020a). Seven genetic clades (S, L, O, V, G, GR, and GH) have been described over time that are spread throughout different countries (Alm et al. 2020). These clades represent a challenge for public health as re-infection cases with different clade strains have been reported (Grupta et al. 2020; To et al. 2020; Van Elslande et al. 2020). In fact, more than 200 candidates for vaccines against SARS-CoV-2 and several antivirals are already being developed (Hu et al. 2020; WHO, 2020b). Most of vaccines and therapeutic drugs are directed towards the spike glycoprotein (S) that is responsible for entering the host cell through recognition of the receptor ACE2 with the receptor binding protein (RBD) (Alexpandi et al. 2020; Olaleye et al. 2020; Shang et al. 2020; Trezza et al. 2020; WHO, 2020b). Even though some studies have determined the nucleotide evolutionary rate of SARS-CoV-2 using the entire genome (Giovanetti et al. 2020; van Dorp et al. 2020), those values are slower and do not represent the real mutation capacity of the S region alone. Only one study has reported the nucleotide evolutionary rate of the S genomic region in the first four months of the pandemic, but without differentiating the seven viral clades, which can be relevant in therapeutics and re-infections (Pereson et al. 2020). Thus, the aim of this study was to determine the nucleotide evolutionary rate and the haplotype network of the S region for SARS-CoV-2 in general and for each of the seven genetic clades during the first nine months of pandemic.

## 2. Materials and Methods

### 2.1 Datasets

In order to generate datasets representing different geographic regions and time evolution for each of the seven clades of SARS-CoV-2, from December 2019 to September 2020, data of complete genome sequences available at GISAID (https://www.gisaid.org/) on September 2020 were randomly monthly collected for several geographic regions. Data inclusion criteria were: a.-complete genomes, b.-high coverage level, and c.-human host only (no other animals or environmental samples). Complete genomes were aligned using MAFFT against the Wuhan-Hu-1 reference genome (NC_045512.2, EPI_ISL_402125). The resulting multiple sequence alignments were split in a dataset corresponding to the S region [3,822nt (21,563-25,384)] and RBD (included in S) [762nt (22,550-23,311)].

### 2.2 Phylogenetic and statically analysis / Genetic characterization

Patterns of genetic diversifications for both genomic regions S and RBD for each clade were analyzed using the median-joining reconstruction method at the PopART v1.7.2 software (Leigh & Bryant, 2015). Haplotypes shared among all clades were analyzed in Arlequin 3.5.2.2 software (Excoffier & Lischer, 2010). Polymorphism indices were calculated separately for each clade with DnaSPv. 6.12.01 (Rozas et al. 2017).

### 2.3 Nucleotide evolutionary rate

The estimation of the nucleotide evolutionary rate for the entire S-coding region datasets were carried out with the Beast v1.8.4 program package (Suchard et al. 2018) at the CIPRES Science Gateway server (Miller et al. 2010). The temporal calibration was established by the samples’ date of sampling. The best nucleotide substitution model was selected according to the Bayesian information criterion (BIC) method in IQ-TREE v1.6.12 software (Kalyaanamoorthy et al. 2017). The analysis was performed under a relaxed (uncorrelated lognormal) molecular clock model recommended previously by Duchene & col., (Duchene et al. 2020) with an exponential demographic model (Grassly & Fraser, 2008) ^[23]^. Analyses were run for 8×10^6^ generations and sampled every 8×10^5^ steps. The convergence of the “meanRate” and “allMus” parameters [effective sample size (ESS) ≥ 200, burn-in 10%] was verified with Tracer v1.7.1 (Rambaut et al. 2018). The obtained substitution rate was probed against 10 independent replicates of the analysis with the time calibration information (date of sampling) randomized as described by Rieux & Khatchikian, 2017 (Rieux & Khatchikian, 2017).

## 3. Results

### 3.1 Datasets

Three-hundred sequences were randomly selected for each clade. Two thousand and one hundred sequences were curated and selected for the analysis. Table 1 shows the SARS-CoV-2 sequences included for every month and clade.

**Table 1.**
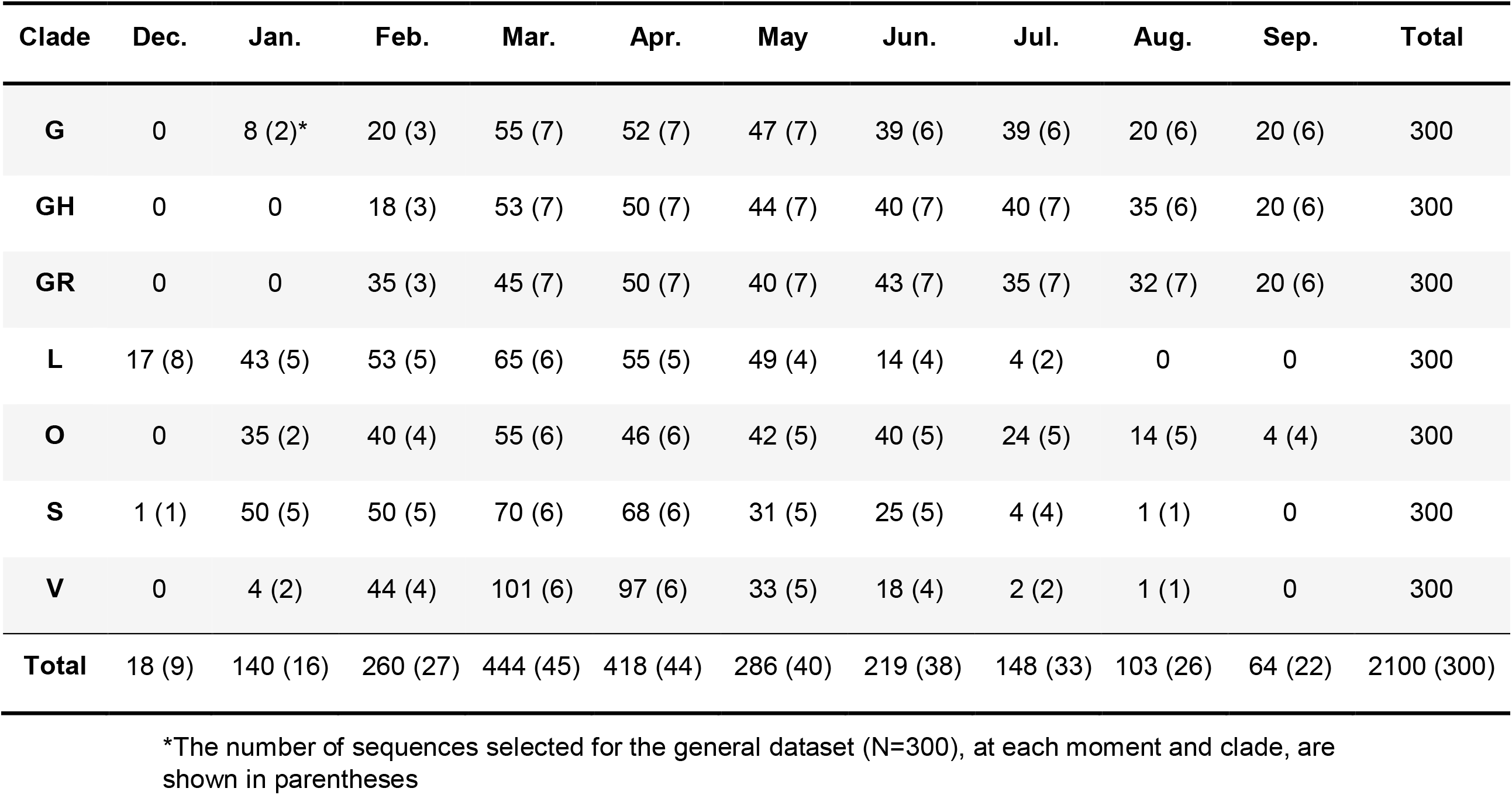
Number of SARS-CoV-2 sequences from GISAID database on September 18^th^, by month and clade as per the selection criteria (Temporal structure).

| Clade | Dec. | Jan. | Feb. | Mar. | Apr. | May | Jun. | Jul. | Aug. | Sep. | Total |
| --- | --- | --- | --- | --- | --- | --- | --- | --- | --- | --- | --- |
| **G** | 0 | 8 (2)* | 20 (3) | 55 (7) | 52 (7) | 47 (7) | 39 (6) | 39 (6) | 20 (6) | 20 (6) | 300 |
| **GH** | 0 | 0 | 18 (3) | 53 (7) | 50 (7) | 44 (7) | 40 (7) | 40 (7) | 35 (6) | 20 (6) | 300 |
| **GR** | 0 | 0 | 35 (3) | 45 (7) | 50 (7) | 40 (7) | 43 (7) | 35 (7) | 32 (7) | 20 (6) | 300 |
| **L** | 17 (8) | 43 (5) | 53 (5) | 65 (6) | 55 (5) | 49 (4) | 14 (4) | 4 (2) | 0 | 0 | 300 |
| **O** | 0 | 35 (2) | 40 (4) | 55 (6) | 46 (6) | 42 (5) | 40 (5) | 24 (5) | 14 (5) | 4 (4) | 300 |
| **S** | 1 (1) | 50 (5) | 50 (5) | 70 (6) | 68 (6) | 31 (5) | 25 (5) | 4 (4) | 1 (1) | 0 | 300 |
| **V** | 0 | 4 (2) | 44 (4) | 101 (6) | 97 (6) | 33 (5) | 18 (4) | 2 (2) | 1 (1) | 0 | 300 |
| **Total** | 18 (9) | 140 (16) | 260 (27) | 444 (45) | 418 (44) | 286 (40) | 219 (38) | 148 (33) | 103 (26) | 64 (22) | 2100 (300) |

\*The number of sequences selected for the general dataset (N=300), at each moment and clade, are shown in parentheses

### 3.2 Phylogenetic and statically analysis / Genetic characterization

The haplotype networks (Figure 1) reflect the diversity indices results as a star shape with multiple haplotypes with few nucleotide differences that diverge from a common ancestor. In all cases, the RBD diversification is lower than the spike one, being the lowest for clades S and V. For the S-coding region, 479 different haplotypes were defined in the seven analyzed clades. The number of haplotypes observed among clades ranged from 53 for the V clade to 89 for the GH and GR clades (Table 2). The major haplotype 1 (Hap-1), defined by amino acids S12, L18, R21, A222, N439, S477, T478, A522, E583, G614, Q675, E780, D936, V1068, and P1263 was the most frequent for clades G (54%), GH (54%), and GR (56%). However, other 10 haplotypes with amino acid changes respect to the Hap-1 were also observed. On the other hand, haplotype 252 (Hap-252), defined by amino acids L5, L8, H49, V367, A575, D614, A829, A846, D1084, and A1087 was the most frequent for clades L (63.3%), O (39.7%), S (51.7%), and V (70%). In addition, other 10 haplotypes showed one amino acid change respect to Hap-252. Table 3 shows the frequency of each haplotype with amino acid changes. The haplotype diversity was moderate to high in every clade, ranging from Hd = 0.507 to 0.793 (Table 2). In contrast, nucleotide diversity was relatively low for each clade, ranging between π = 0.0018 for V and π = 0.0040 for O (Table 2). Although overall diversity was similar among different clades, haplotype and nucleotide diversities were both the lowest for V. On the other hand, haplotype and nucleotide diversity was higher for G, GH, GR, and O (Table 2). The RBD region showed indices with a similar trend but with lower values compared to the S region.

**Table 2.** Summary of the haplotype and nucleotide diversity indices for the entire Spike and the Receptor Binding-Domain coding regions for each clade of SARS-COV2.

| ***SPIKE*** |  |  |  |  |
| --- | --- | --- | --- | --- |
| **Clade** | **S** | **H** | **Hd** | **π** |
| **G** | 100 | 86 | 0.704 ± 0.030 | 0.00037 ± 0.00003 |
| **GH** | 102 | 89 | 0.704 ± 0.030 | 0.00038 ± 0.00003 |
| **GR** | 112 | 89 | 0.683 ± 0.031 | 0.00038 ± 0.00003 |
| **L** | 87 | 76 | 0.598 ± 0.035 | 0.00023 ± 0.00002 |
| **O** | 81 | 68 | 0.793 ± 0.019 | 0.00040 ± 0.00002 |
| **S** | 72 | 60 | 0.716 ± 0.027 | 0.00031 ± 0.00002 |
| **V** | 56 | 53 | 0.507 ± 0.036 | 0.00018 ± 0.00002 |
| **General** | 134 | 107 | 0.857 ± 0.015 | 0.00052 ± 0.00003 |
| ***RBD*** |  |  |  |  |
| **Clade** | **S** | **H** | **Hd** | **π** |
| **G** | 15 | 15 | 0.183 ± 0.030 | 0.00028 ± 0.00005 |
| **GH** | 17 | 19 | 0.196 ± 0.031 | 0.00032 ± 0.00006 |
| **GR** | 23 | 23 | 0.281 ± 0.034 | 0.00041 ± 0.00006 |
| **L** | 15 | 14 | 0.104 ± 0.024 | 0.00016 ± 0.00004 |
| **O** | 9 | 9 | 0.193 ± 0.030 | 0.00027 ± 0.00004 |
| **S** | 3 | 4 | 0.027 ± 0.013 | 0.00003 ± 0.00002 |
| **V** | 3 | 4 | 0.020 ± 0.011 | 0.00003 ± 0.00001 |
| **General** | 17 | 17 | 0.166 ± 0.029 | 0.00026 ± 0.00005 |

S= Number of variable sites; H = Number of haplotypes; Hd = Haplotype diversity; π = Nucleotide diversity (per site)

**Table 3.** Frequency of haplotypes with amino acid changes in the spike for each clade of SARS-COV2.

| Clade/<br>Haplotype (aa Change<br>respect to Hap-1) | G<br>N (%) | GH<br>N (%) | O<br>N (%) | GR<br>N (%) | Clade/<br>Haplotype (aa Change<br>respect to Hap-252) | L<br>N (%) | O<br>N (%) | S<br>N (%) | V<br>N (%) |
| --- | --- | --- | --- | --- | --- | --- | --- | --- | --- |
| Hap-1 (*) | 162 (54) | 162 (54) | 25 (8.3) | 168 (56) | Hap-252 (#) | 190 (63.3) | 119 (39.7) | 154 (51.3) | 210 (70) |
| Hap-7 (S477N) | 5 (1.7) |  |  | 10 (3.4) | Hap-254 (H49Y) | 3 (1) |  |  |  |
| Hap-34 (N439K) | 4 (1.3) |  |  |  | Hap-256 (D1084Y) | 3 (1) |  |  |  |
| Hap-67 (P1263L) | 4 (1.3) |  |  |  | Hap-282 (NC) | 6 (2) | 62 (20.7) |  |  |
| Hap-86 (L18F, A222V) | 20 (6.8) |  |  |  | Hap-291 (L5F) |  |  |  | 10 (3.3) |
| Hap-90 (A522S, E780C) |  | 5 (1.7) |  |  | Hap-320 (A575S) | 8 (2.6) |  |  |  |
| Hap-91 (E780C) |  | 6 (2) |  |  | Hap-324 (A1087S) | 3 (1) |  |  |  |
| Hap-105 (D936Y) |  | 10 (3.4) |  |  | Hap-367 (L8V) |  |  | 17 (5.7) |  |
| Hap-137 (E583D) |  | 3 (1) |  |  | Hap-382 (V367F) |  | 5 (1.7) |  |  |
| Hap-171 (Q675R) |  |  |  | 3 (1) | Hap-384 (D614A) |  |  | 3 (1) |  |
| Hap-187 (S12F) |  |  |  | 3 (1) | Hap-415 (A829T) |  |  | 5 (1.7) |  |
| Hap-226 (T478I) |  |  |  | 8 (2.6) | Hap-437 (A846S) |  |  |  |  |
| **Total** | **300 (100)** | **300 (100)** |  | **300 (100)** | **Total** | **300 (100)** | **300 (100)** | **300 (100)** | **300 (100)** |

2 N: Number, Hap= haplotype, aa= Amino acid, NC= No amino acid changes
\* Hap-1: S12, L18, R21, A222, N439, S477, T478, A522, E583, G614, Q675, E780, D936, V1068, and P1263.

**Figure 1.**
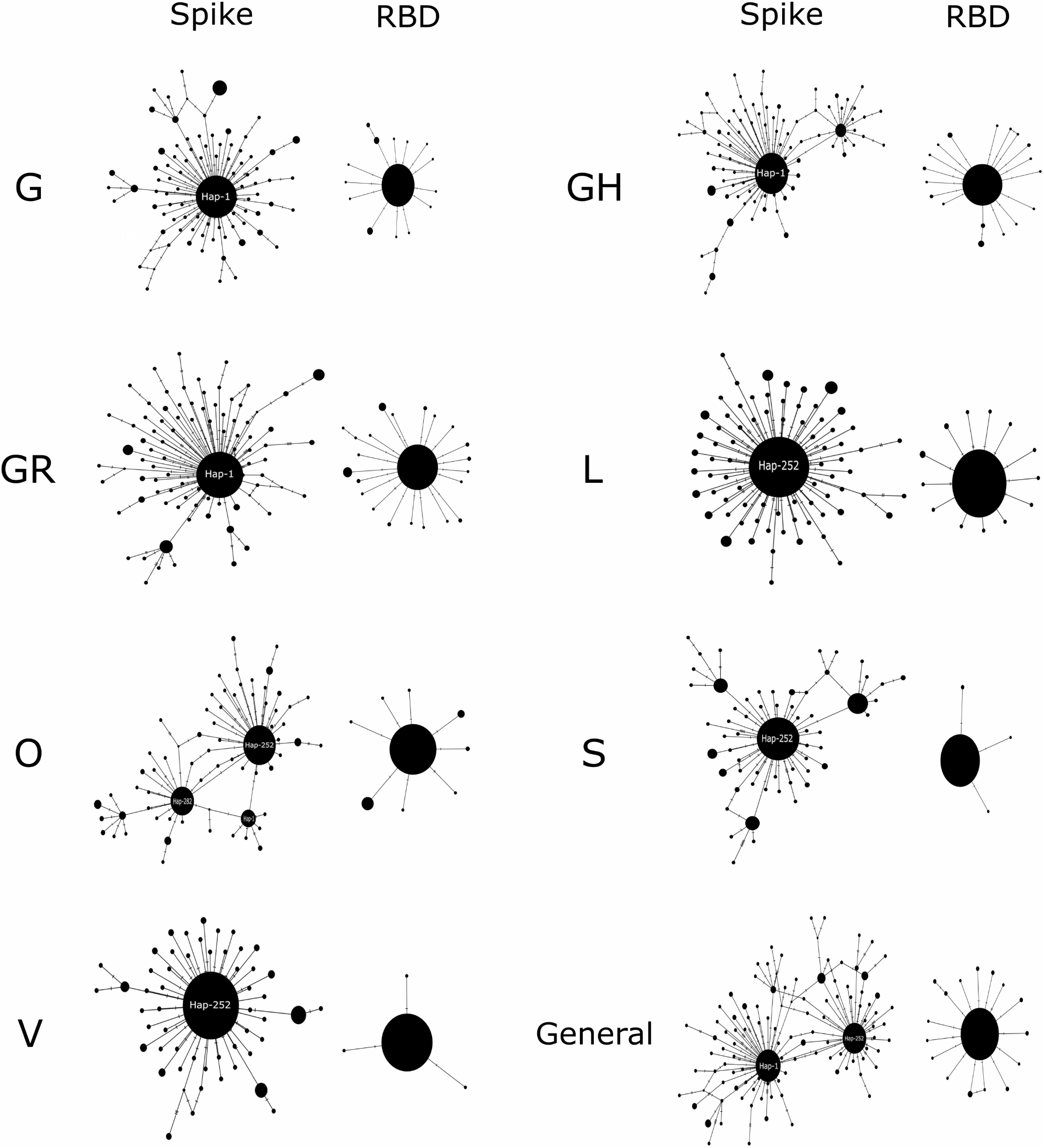
Median-joining haplotype networks. The seven clades of SARS-CoV 2 described to date are compared to both the entire Spike and the RBD coding region. The diameters of the spheres are proportional to the frequency of haplotypes. The main haplogroups are indicated.

### 3.3 Evolutionary rate

After nine months of pandemic, the estimated evolutionary rate for the S genomic region of SARS-CoV-2 was 1.08 x 10^−3^ nucleotide substitutions per site per year (s/s/y) (95% HPD interval 7.94 x 10^−4^ to 1.41 x 10^−3^ s/s/y). Additionally, the nucleotide evolutionary rate for the different genetic clades ranged between 1.06 x 10^−3^ and 1.69 x10^−3^ s/s/y (Table 4). A date-randomization analyses showed no overlapping between the 95% HPD substitution-rate intervals obtained from real data and from date-randomized datasets for all clades (Figure 2). Dataset for the clade L did not reach convergence (ESS<200). To verify the reliability of the result, 10 independent runs were performed. All of them converged in a similar posterior distribution. Likewise, for many of the random sample datasets, convergence was not achieved (ESS between 100 and 200). For those datasets that did not reach convergence, two independent runs were carried out and concatenated (Lemey et al. 2009). When the evolutionary rate was analyzed according to the emergence of each clade, founding clades (L, O, S, and V) tended to present evolutionary rates slightly slower than the more recent clades (G, GH, and GR), (p = 0.157).

**Table 4.**
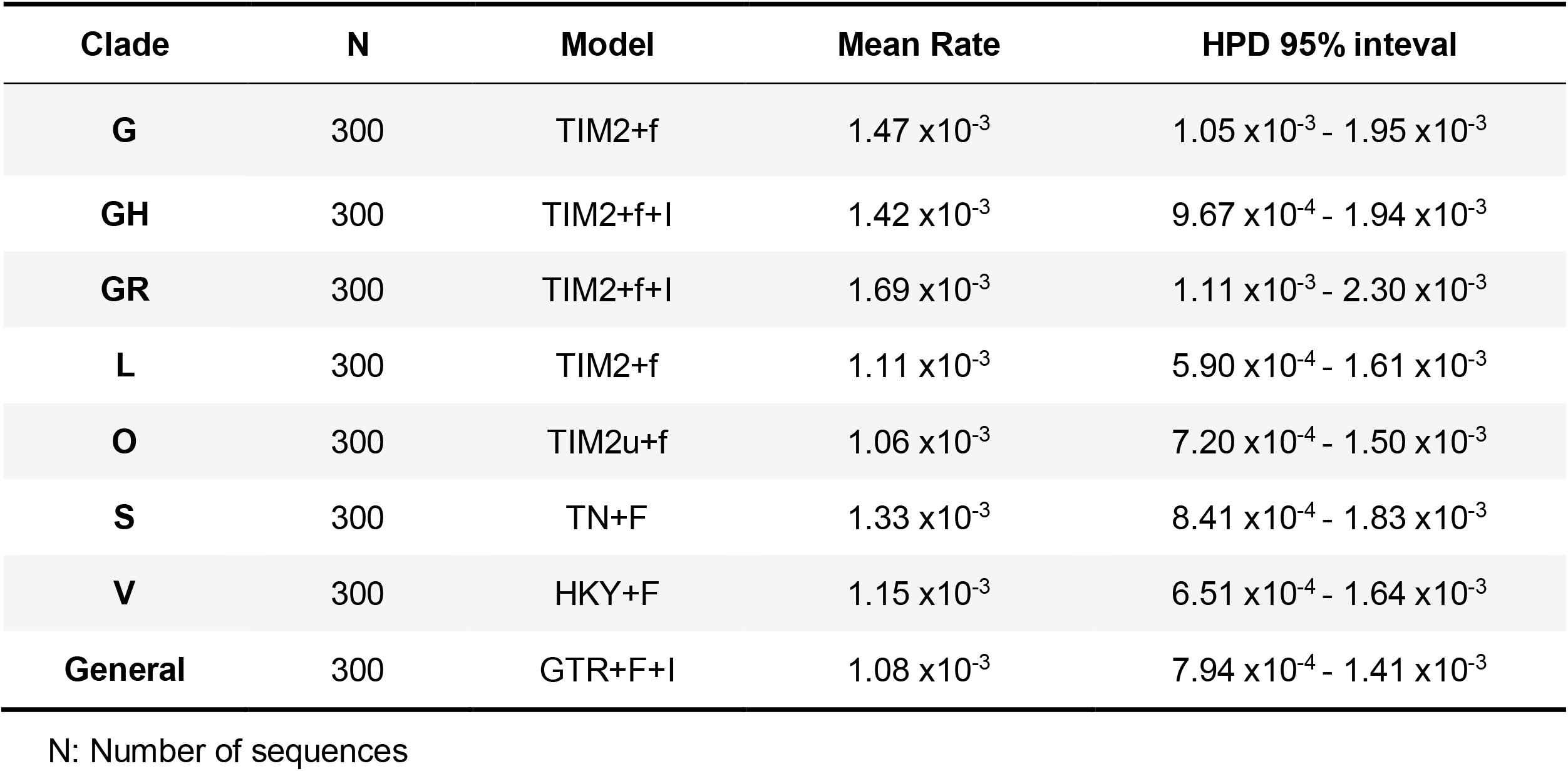
Mean rates of the Spike-coding region (nt = 3822) for each clade of SARS-COV2.

| Clade | N | Model | Mean Rate | HPD 95% interval |
| --- | --- | --- | --- | --- |
| **G** | 300 | TIM2+f | $1.47 \times 10^{-3}$ | $1.05 \times 10^{-3} - 1.95 \times 10^{-3}$ |
| **GH** | 300 | TIM2+f+I | $1.42 \times 10^{-3}$ | $9.67 \times 10^{-4} - 1.94 \times 10^{-3}$ |
| **GR** | 300 | TIM2+f+I | $1.69 \times 10^{-3}$ | $1.11 \times 10^{-3} - 2.30 \times 10^{-3}$ |
| **L** | 300 | TIM2+f | $1.11 \times 10^{-3}$ | $5.90 \times 10^{-4} - 1.61 \times 10^{-3}$ |
| **O** | 300 | TIM2u+f | $1.06 \times 10^{-3}$ | $7.20 \times 10^{-4} - 1.50 \times 10^{-3}$ |
| **S** | 300 | TN+F | $1.33 \times 10^{-3}$ | $8.41 \times 10^{-4} - 1.83 \times 10^{-3}$ |
| **V** | 300 | HKY+F | $1.15 \times 10^{-3}$ | $6.51 \times 10^{-4} - 1.64 \times 10^{-3}$ |
| **General** | 300 | GTR+F+I | $1.08 \times 10^{-3}$ | $7.94 \times 10^{-4} - 1.41 \times 10^{-3}$ |

N: Number of sequences

**Figure 2.**
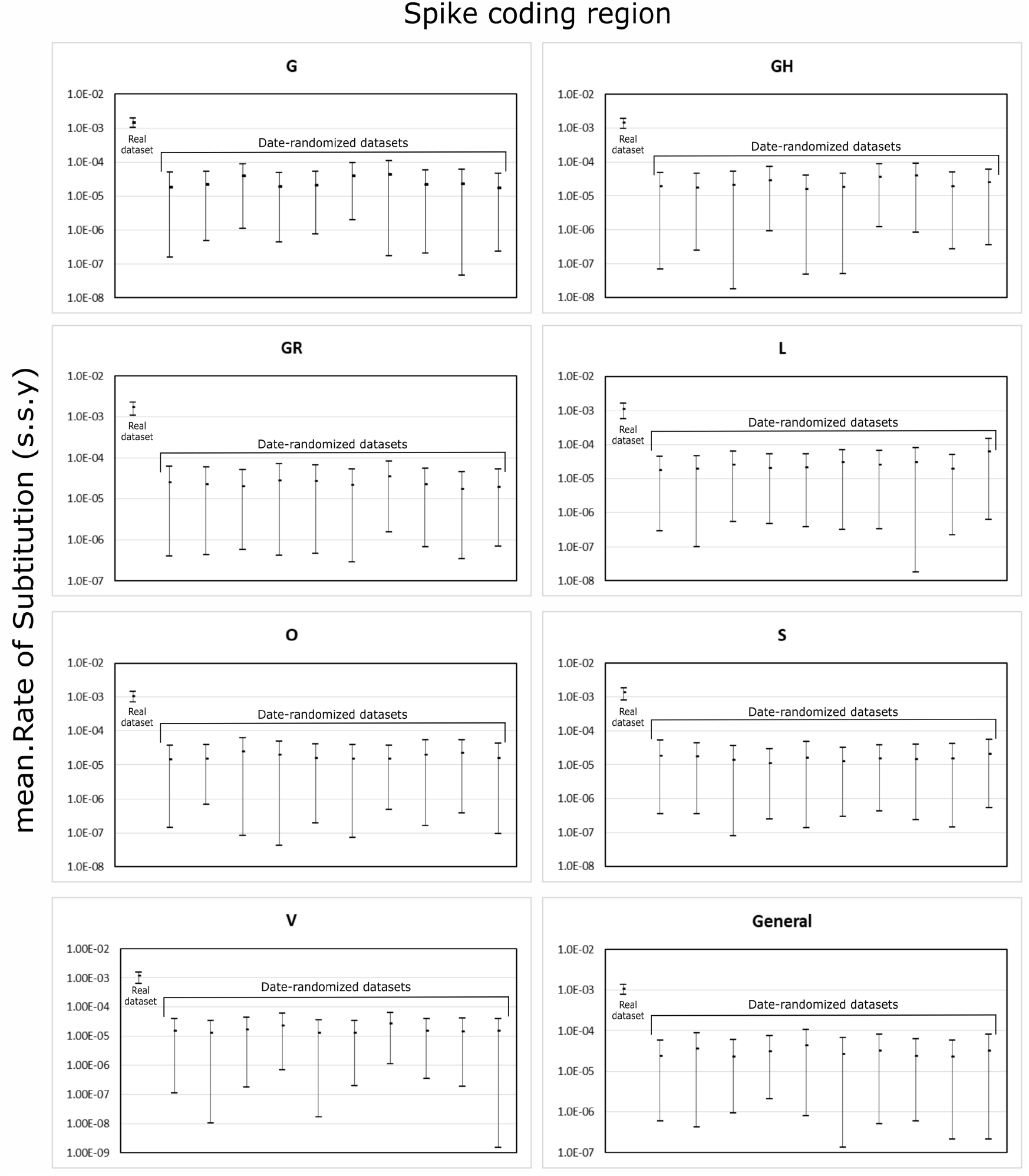
Test of temporal structure. Comparison of the evolutionary rates estimated for the original dataset vs. the date-randomized ones. This analysis was performed for the Spike-coding region (3822nt) of each clade. s.s.y = substitutions/site/year.

## 4. Discussion

The evolutionary characterization of spike genomic region of SARS-CoV-2 is crucial to estimate the course that re-infections, vaccines, and therapeutics would have in the pandemic’s future. In this study, the evolutionary rate of the most important SARS-CoV-2 protein for vaccine development was estimated in general and separately for each genetic clade described in GISAID. In this context, the spike haplotype network showed a founding central paternal haplogroups from which multiple sequences with modest changes derived. Overall, the nucleotide evolutionary rate after nine months of pandemic was similar for each clade.

At the beginning of the pandemic, the most prevalent clades were L, O, V, and S. Later, with the appearance of the D614G mutation in the S protein, clade G emerged and remained with a high and stable prevalence. After this initial step, the GR clade has emerged and grown until it became the most prevalent. Finally, the GH clade peaked at 30% in May 2020 and then began to decrease (Alm et al. 2020). In this sense, it is important to highlight that clades with the mutation D614G in the S protein (clades G, GH, and GR) have been suggested to present a higher transmission efficiency although they would not be associated with a more severe pathogenesis (Korber et al. 2020).

Therefore, in order to describe the evolution of the S protein variants, the study of haplotypes network in all seven clades and for both regions (S and RBD alone) was performed. This analysis showed several identical sequences grouped together resulting in a star-shaped network, which is characteristic of viral outbreaks (Liu et al. 2020). For the spike, this general analysis was supported by statistics that show a large number of haplotypes with a small number of nucleotide changes (low nucleotide diversity). However, for the RBD region, an increase in identical haplotypes was observed, which translates into a decrease in other parameters (H, Hd, and Π). This may be due to the conserved nature of the cell receptor-binding region, which is necessary for the infection of target cells. It is noteworthy that the lowest gene and nucleotide diversities observed for clade V, in both S and RBD, could be the result of fewer sequences available for this clade during the nine months analyzed here. In this way, it can be observed that more than 90% of the V clade sequences were distributed in four months (February to May). On the other hand, the highest nucleotide diversity observed in clade O is the result of a less clearly defined pattern of mutations (Mercatelli & Giorgi, 2020). Several amino acid changes detected in the haplotypes present in our analysis are part of the RBD (V367F, S477N, N439K, T478I, and A522S). From these amino acid changes, positions 367 and 439 were associated with the binding affinity of RBD (Teng et al. 2020; Yi et al. 2020). Additionally, the mutation L5F in the signal peptide was present in 3.3% of members belonging clade V (Korber et al. 2020). Other changes associated to relevant functions (Korber et al. 2020; Teng et al. 2020) such as H49Y in clade L (associated with monomer stability), A829T in clade S (fusion peptide), D936Y in clade GH [Heptad repeat 1 (HR1) associated with monomer stability], and P1263 in clade G (present in the cytoplasmic tail), were also detected in 1% to 3.4%.

The evolutionary characterization of the wide spectrum of haplotypes contributes to determine the haplotype significance and its association with disease severity, response to antivirals, development of vaccines, and host genetic factors.

The evolutionary rate of S protein estimated for all together clades was significantly higher than that previously reported by analyzing the entire genome (van Dorp et al. 2020; Liu et al. 2020). This is expected since the complete genome includes several genomic regions with a high degree of conservation, while the S region is one of the most rapidly evolving in the SARS-CoV-2 genome (Pereson et al. 2020). Nonetheless, the spike evolution rate was quite similar to that obtained by analyzing this region during the first four months of the pandemic (Pereson et al. 2020). Although the evolutionary rate of all clades was similar, the founding clades (L, O, V, and S) showed evolutionary rates slightly lower than the most recent and currently more distributed ones (G, GH, and GR). This could be endorsed to the spread process in human populations since they are the most widely disseminated clades around the world.

This study provides substantial data on the evolutionary process of S protein in the different clades of a virus that infects a susceptible population where a massive active immunization process has not yet been carried out. However, as it was aforementioned, the evolutionary rate of the S region remained stable throughout the nine considered months. In coming months, this scenario may modify and it would be necessary to re-evaluate the results from this study. In fact, a new clade named GV was described last months (Hadfield et al. 2018). The inclusion in the study of only 2,100 of the 73,393 available sequences on September 2020 is a limitation that imply a bias in the obtained results, although the sequence selection process was carefully carried out in order to generate a representative dataset from different time courses and a wide geographic range.

## 5. Conclusions

Since the S protein of SARS-CoV-2 mediates the entry in the host cell and is the target for most therapeutic candidates, it is essential to know the way this genomic region is evolving, given that changes in this protein could have consequences on viral transmission, response to antivirals, and efficacy of vaccines. On this basis, the results obtained in this work about the evolutionary rate of the spike protein during the first nine months of pandemic are very significant. Furthermore, the evolutionary study of each separate clade ads to the virus knowledge and deserves to be assessed in more detail since re-infection by a different phylogenetic clade has been reported.

## Funding

None

## Disclosures

None.

## Author contributions

MJP: Data curation, acquisition of data, analysis and interpretation of data, drafting the article, final approval of the version to be submitted.

DMF: Data curation, Validation, drafting the article, final approval of the version to be submitted.

APM: Data curation, Validation, revising the article critically for important intellectual content, final approval of the version to be submitted.

PB: Data curation, acquisition of data, analysis and interpretation of data, revising the article critically for important intellectual content, final approval of the version to be submitted.

GG: Data curation, acquisition of data, analysis and interpretation of data, drafting the article, final approval of the version to be submitted.

FAD: Conception and design of the study, acquisition of data, analysis and interpretation of data, drafting the article, final approval of the version to be submitted.

## Acknowledgements

MJP, DMF, PB and FAD are members of the National Research Council (CONICET). We would like to thank to the researchers who generated and shared the sequencing data from GISAID (https://www.gisaid.org/) and Mrs. Silvina Heisecke from CEMIC-CONICET for providing language assistance.

## Notes

### Competing Interest Statement

The authors have declared no competing interest.

